# Predicting task-related brain activity from resting-state brain dynamics with fMRI Transformer

**DOI:** 10.1101/2024.05.29.596544

**Authors:** Junbeom Kwon, Jungwoo Seo, Heehwan Wang, Taesup Moon, Shinjae Yoo, Jiook Cha

## Abstract

Accurate prediction of the brain’s task reactivity from resting-state functional magnetic resonance imaging (fMRI) data remains a significant challenge in neuroscience. Traditional statistical approaches often fail to capture the complex, nonlinear spatiotemporal patterns of brain function. This study introduces SwiFUN (Swin fMRI UNet Transformer), a novel deep-learning framework designed to predict 3D task activation maps directly from resting-state fMRI scans. SwiFUN leverages advanced techniques such as shifted window-based self-attention, which helps to understand complex patterns by focusing on varying parts of the data sequentially, and a contrastive learning strategy to better capture individual differences among subjects. When applied to predicting emotion-related task activation in adults (UK Biobank, n=7,038) and children (ABCD, n=4,944), SwiFUN consistently achieved higher overall prediction accuracy than existing methods across all contrasts; it demonstrated an improvement of up to 27% for the FACES-PLACES contrast in ABCD data. The resulting task activation maps revealed individual differences across cortical and subcortical regions associated with sex, age, and depressive symptoms. This scalable, transformer-based approach potentially reduces the need for task-based fMRI in clinical settings, marking a promising direction for future neuroscience and clinical research that enhances our ability to understand and predict brain function.

## 1. Introduction

Task-based functional Magnetic Resonance Imaging (fMRI) has been instrumental in cognitive neuroscience, offering insights into the functional neuroanatomy associated with adaptive and maladaptive cognition and behavior. This method holds promise for predicting individual cognitive functions and psychological disorders (Brodersen et al., 2011; Frässle et al., 2020), often outperforming resting-state fMRI in accuracy (Gal, Coldham, et al., 2022; Gal, Tik, et al., 2022; Greene et al., 2018; Sripada et al., 2020). Its utility extends to pre-surgical planning, where it aids in identifying functionally impaired brain regions, thus helping to reduce the risk of cognitive impairments post-surgery (Niu et al., 2021; Parker Jones et al., 2017). However, implementing task-based functional magnetic resonance imaging (fMRI) in practical settings like clinical practice is challenging. This is due to issues such as ensuring participant compliance and motivation, as well as the necessity for strict experimental control. These issues become particularly pronounced in specific groups such as children, the elderly, and individuals with neurocognitive disorders or severe psychiatric conditions (Bernstein-Eliav & Tavor, 2022; Zhang et al., 2021).

Recent research is steering towards a significant shift in predicting task reactivity, focusing on resting-state fMRI, which reveals the brain’s intrinsic activity patterns (Bernstein-Eliav & Tavor, 2022). This approach is based on the premise and findings of close relationships between resting-state and active-state functional networks (Cole et al., 2014; Elliott et al., 2019; Smith et al., 2009), suggesting that the ”connectivity fingerprint” derived from resting-state fMRI reflects salient individual differences in cognitive functioning (Finn et al., 2015; Ito et al., 2022; Schultz et al., 2022). Such fingerprints have been shown to predict task activation across various cognitive tasks (Cohen et al., 2020; Cole et al., 2016; Tavor et al., 2016; Tripathi & Somers, 2023; Zheng et al., 2022), even in patients with neurological disorders (Niu et al., 2021; Parker Jones et al., 2017; Tik et al., 2021). The consistent correlation between resting-state functional connectivity and task activation was validated across diverse sites, MRI vendors, and age groups in multiple cognitive tasks (Tik et al., 2023). In addition, task activation maps derived by resting-state functional connectivity achieved more accurate predictions of cognitive functioning than resting-state functional connectivity alone (Gal, Tik, et al., 2022) Furthermore, researchers can investigate underlying functional connections of task-related cognitive processes by identifying resting-state functional connectivity predictive of task-induced brain activity (Izakson et al., 2023; Tik et al., 2021).

The advent of deep learning marks a significant evolution in connectivity fingerprinting from fMRI data, enabling enhanced predictive accuracy and reliability over traditional methods. BrainSurfCNN, a surface-based deep convolutional neural network pre-trained on extensive datasets and fine-tuned for specific applications, has shown marked capability in predicting task activation maps from resting-state gray-ordinate fMRI data (Ngo et al., 2022; M. Nguyen et al., 2023). These models surpass traditional approaches that rely on rule-based features of the functional patterns (e.g., independent component analysis followed by dual regression (Tavor et al., 2016), or stochastic probabilistic functional modes (Zheng et al., 2022)), which only partially capture the brain’s spatial and temporal dynamics. Other recent studies suggested that applying deep learning models directly to volumetric fMRI data can better capture subtle individual differences in fMRI data than deep learning models relying on handcrafted features, showing outstanding predictive performance for cognitive and biological variables (Kim et al., 2024; Malkiel et al., 2022; S. Nguyen et al., 2020; Rosenman et al., 2023). The question then arises: Can we predict task-related activations from resting-state brain activity by capturing spatiotemporal patterns directly from fMRI data?

In response to this challenge, we introduce SwiFUN (Swin fMRI Transformer with UNET), a pioneering end-to-end deep learning model designed to generate task activation maps from resting-state fMRI data. SwiFUN, leveraging the innovative Swin (shifted window) UNETR architecture (Hatamizadeh et al., 2022) and a Swin 4D fMRI Transformer (Kim et al., 2024) combined with contrastive learning strategy (Ngo et al., 2022), represents the first application of such technology in human neuroimaging studies. Our findings suggest that SwiFUN can learn rich spatiotemporal representations from resting-state fMRI data, significantly improving the prediction of human brain activity during specific tasks. This novel approach may potentially transform neuroscience research, offering a more effective and inclusive method of exploring the functional neuroanatomy of cognition and behavior.

## 2. Methods

### 2.1 Experimental Setup

#### 2.1.1. Data

We used 3-Tesla resting-state functional Magnetic Resonance Imaging (rsfMRI) and task-based fMRI (tfMRI) data from participants in the UK Biobank and the Adolescent Brain Cognitive Development study. In contrast to the conventional approach using rsfMRI data and tfMRI in surface space (CIFTI: (’grayordinate’ surface + volume)) (Van Essen et al., 2013), we used minimally preprocessed resting-state fMRI data and volumetric z-score task activation maps related to specific tasks or visual stimuli. We masked out the irrelevant brain and non-brain voxels using two brain atlas images to restrict the analysis to the comparable brain regions as the ConnTask, a machine learning model depending on parcellation (Gal, Coldham, et al., 2022; Gal, Tik, et al., 2022; Tavor et al., 2016; Tik et al., 2021, 2023). Specifically, we employed 100 cortical parcels defined by Schaefer et al., 2018, which are assigned to one of the seven brain networks and widely used in the previous research (Cohen et al., 2020; Gal, Coldham, et al., 2022; Gal, Tik, et al., 2022; Tavor et al., 2016), and Harvard-Oxford cortical and subcortical structural atlases for evaluating the model’s performance in predicting subcortical brain activity (Desikan et al., 2006; Frazier et al., 2005; Goldstein et al., 2007; Makris et al., 2006). As a result, a total of 132,032 (Schaefer) and 146,025 (Harvard-oxford) valid voxels in task activation maps and each resting-state fMRI volume were selected for our analysis.

##### UK Biobank data

UK Biobank (UKB) is a large biomedical database that contains health-related information from half a million UK participants. To evaluate the model’s ability to generate task activation maps, we ran the analysis on the preprocessed resting-state and task-state fMRI of 7,038 individuals (age = 54.971 ± 7.53 years, 52.7% female) from UK Biobank release 2. The detailed acquisition protocol and preprocessing process are described in Miller et al., 2016 and Alfaro-Almagro et al., 2018. The fMRI data has a resolution of 2.4 * 2.4 * 2.4 mm, having TR of 0.735 s and TE of 39ms. The scan duration is 6 minutes (490 time points) for resting-state fMRI and 4 minutes (332 time points) for task fMRI. The initial preprocessing performed by the UKB Brain imaging team includes motion correction, group-mean intensity normalization, highpass temporal filtering, and EPI warping. Spatial smoothing with a Gaussian kernel of FWHM 5mm was applied to task fMRI before intensity normalization. Unlike task fMRI, the resulting resting-state fMRI scans were further ICA+FIX cleaned to remove structured artifacts (Beckmann & Smith, 2004; Salimi-Khorshidi et al., 2014). The task used is the Hariri faces/shapes ”emotion” task, where participants viewed faces or shapes sequentially in each block of trials. Z-statistics of three contrasts were estimated from task fMRI using FSL FEAT (Woolrich et al., 2004): Shapes, Faces, and Faces-Shapes (Barch et al., 2013). The resting-state fMRI scans and task contrast maps were then registered to standard MNI space (Grabner et al., 2006).

##### Adolescent Brain Cognitive Development data

Adolescent Brain Cognitive Development (ABCD) study is the largest longitudinal study of brain and cognitive development in the United States. We used resting-state and task-state fMRI scans during emotional n-back (EN-back) task from 4,944 adolescents (age = 9.95 ± 0.63 years, 49.7% female) (Casey et al., 2018) from release 2. The fMRI data has a resolution of 2.4 * 2.4 * 2.4 mm, having TR of 0.8 s and TE of 30ms. The duration of the scan amounts to 5 minutes (383 time points) for resting-state fMRI and 4.8 minutes (362 time points) for task fMRI. Using fMRIprep (Esteban et al., 2019), an automated preprocessing pipeline for structural and functional MRI, we conducted brain extraction, slice time correction, and confounds estimation. Then, we spatially normalized fMRI data to the standard MNI space for the pediatric brains (i.e., MNIPediatricAsym space;(V. Fonov et al., 2011; V. S. Fonov et al., 2009)). For resting-state fMRI, we additionally applied low pass filtering, head movement correction, and artifact removal, regressing out signals from non-grey matters (aCompcor). We applied spatial smoothing with a Gaussian kernel with FWHM of 5mm to task fMRI and normalized the signal of each voxel by the mean across the time of each voxel following Chaarani et al., 2021.

We used a generalized linear model (GLM) to estimate the participants’ task-related brain activation map in the Emotional N-back task. We created a task model considering a block design following Chaarani et al., 2021, with a task block duration of 24.5 seconds and a fixation block duration of 14 seconds. We used a total of nine conditions of blocks as GLM regressors (i.e., block of 0-back positive face, 0-back negative face, 0-back neutral face, 0-back place, 2-back positive face, 2-back negative face, 2-back neutral face, 2-back place, and fixation). We used a total of 15 nuisance regressors for GLM, including confounding variables calculated by fMRIPrep, such as average signal within brain mask, white matter mask, and CSF mask, as well as six motion parameters and their time derivatives. We used SPM’s canonical hemodynamic response function (double-gamma SPM model with time derivative). To remove low-frequency fluctuations in the BOLD time series, we used high pass filtering with a Discrete Cosine Transform (DCT) basis (cut-off freq = 1/128 Hz) as the drift model. We censored frames with framewise displacement (FD) of 0.9mm or higher. We estimated four z-statistics maps for the following contrasts: Places, Faces, Faces-Places, and 2back-0back.

##### Feature extraction for resting-state functional modes

Previous studies have used resting-state functional modes for predicting task activation maps (Gal, Coldham, et al., 2022; Gal, Tik, et al., 2022; Tavor et al., 2016; Tik et al., 2021, 2023; Zheng et al., 2022). Functional modes refer to consistent spatial patterns of brain activity observed across different individuals. Group-Independent Component Analysis (ICA) is widely used to extract the group-level functional modes from resting-state fMRI. The spatial maps from group-level ICA are data-driven parcellation that extracts independent components (IC) from fMRI data, each representing distinct brain functional networks. UK Biobank executed the group ICA, and UK Biobank provided two versions of group-level ICs (25 and 100) (refer to Miller et al., 2016 for the detailed process). The group-level parcellation results in UK Biobank are available at the following URL: http://biobank.ctsu.ox.ac.uk/crystal/refer.cgi?id=9028. We filtered out components thought to be artefactual from the initial 25 and 100 group-ICA components, leaving 21 and 55 independent components for the dual regression (Miller et al., 2016).

To obtain the group-level functional modes of ABCD data, we carried out group-ICA on the resting-state fMRI of ABCD data. Group ICA was performed only on the specific training subjects, and we excluded them from the rest-to-task fMRI activation map prediction. Only genetically unrelated European subjects with at least 300 resting-state fMRI time points among healthy controls were selected. Healthy controls were defined as subjects with a Child Behavior Checklist (CBCL) total score of 60 or less and a normal KSAD diagnosis based on parent and child questionnaires. This totaled 215 subjects, including 103 males and 112 females. The resting-state fMRI data was spatially smoothed with a 5mm kernel. Then, the group-PCA was applied to the fMRI data by MELODIC’s Incremental Group-PCA, generating 1000 spatial eigenmaps (Smith et al., 2014). The eigenmaps were used to generate group-ICA spatial maps at multiple ICA dimensions by using FSL’s MELODIC function. The ICA dimension used for our experiments was 45.

To derive functional modes for each subject, we performed dual regression using the spatial group IC maps as templates (Nickerson et al., 2017). Before the dual regression, we masked the group IC maps and fMRI scans with a whole-brain mask to exclude unnecessary non-brain voxels. The dual regression consists of two steps. In the first step, the fMRI data were regressed onto the spatial IC maps, resulting in the subject-specific time courses associated with each IC component. In the second step, the previous subject-specific time courses were regressed onto the previous fMRI data, creating individual network-specific spatial IC maps. The individual network-specific spatial IC maps were then used for weighted seed-to-voxel analysis. The individual IC maps were used to regress against the individual resting-state fMRI time series, resulting in a single time series for each spatial map. Subsequently, each time series was correlated with the original fMRI data to generate connectivity maps for each spatial IC. The resulting connectivity features have dimensions of voxels by the number of independent components (ICs).

#### 2.1.2 Measures of Model Performance

We evaluated the predictive performance of task activation maps by assessing, firstly, the overall prediction accuracy and, secondly, how well an individual predicted map identifies individual variability as outlined in previous studies (Tavor et al., 2016; Tik et al., 2021; Zheng et al., 2022). The Pearson correlation coefficient served as our metric for comparing predicted and actual task activation maps.

Our analysis involved calculating pairwise correlation coefficients across N individuals’ actual and predicted activation maps, resulting in an N x N correlation matrix. The diagonal elements, denoted as *C_ii_*, reveal the correlation between predicted and actual activation maps from the same individual (diagonal correlations). Conversely, the off-diagonal elements, *C_ij_* (where *i ≠ j*), represent the correlations between the *i*th actual and *j*th predicted activation maps across different individuals (off-diagonal correlations). We evaluated overall prediction accuracy using the median of diagonal correlations, providing an overview of how well the predicted task activation maps correlated with the actual subject’s maps.

The overall prediction accuracy can be improved by predicting the group mean activation map, which poses a challenge when attempting to predict subtle distinctions in the task activation map among individuals who are performing identical tasks. Therefore, we used three metrics to compare how well each model captures individual differences in task activation maps: the diagonality index, the diagonal percentile mean, and the effect size (*D*) of the Kolmogorov-Smirnov test. The diagonality index, a commonly used metric in previous studies (Tik et al., 2021, 2023; Zheng et al., 2022), is a mean of off-diagonal correlations subtracted from the mean of diagonal correlations. To evaluate whether a task activation map generated by the model for a particular subject specifically predicts the actual activation map for that subject compared to task activation maps generated for other subjects, we developed a new metric, termed the diagonal percentile mean. The diagonal percentile mean is determined by calculating the average percentile of each subject’s diagonal correlation compared to their off-diagonal correlations. The formula for the diagonal percentile mean is as follows:

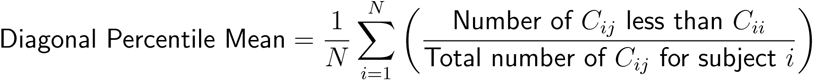

where *C_ij_* is the correlations between the *i*th actual and *j*th predicted activation maps across N different individuals. This metric ranges from 0.5 (indicative of by chance) to 1. If the predicted map for a subject shows lower agreement with the actual map compared to the predicted maps of the others, the diagonal percentile would be closer to 0.5. If there is a high degree of agreement compared to the predicted maps of the others, the diagonal percentile for the subject would be closer to 1. We conducted a Kolmogorov-Smirnov (K-S) test on off-diagonal correlations and diagonal correlations to determine whether there was a significant difference in distributions between the cumulative density functions (CDF) of diagonal and off-diagonal correlations. We used the effect size of the Kolmogorov-Smirnov test (*D*), which represents the maximum difference between the CDFs of the diagonal and off-diagonal correlations, to compare the model’s performances. These metrics altogether may provide a comprehensive evaluation of the predicted activation maps in terms of overall prediction accuracy and individual-level specificity.

### 2.2 Swin fMRI UNet Transformer (SwiFUN)

We implemented a novel deep learning framework called Swin fMRI UNet Transformer (SwiFUN), which can generate task activation maps from spatiotemporal representations. SwiFUN is based on the architecture of the Swin UNet TRansformer (UNETR) model proposed for brain structural segmentation (Hatamizadeh et al., 2022). We adopted the Swin UNETR module from MONAI (Cardoso et al., 2022). As shown in Figure 1, for each contrast, a separate SwiFUN model was trained using a series of fMRI volumes (*T* time points) as input. The goal of each SwiFUN model was to learn the spatiotemporal patterns from the resting-state fMRI data for predicting a single 3D task activation map. The intermediate outputs of each Swin Transformer layer are fed into the UNET decoder through skip connections. This UNET structure enhances training stability and facilitates the generation of higher-resolution image information.

**Figure 1:**
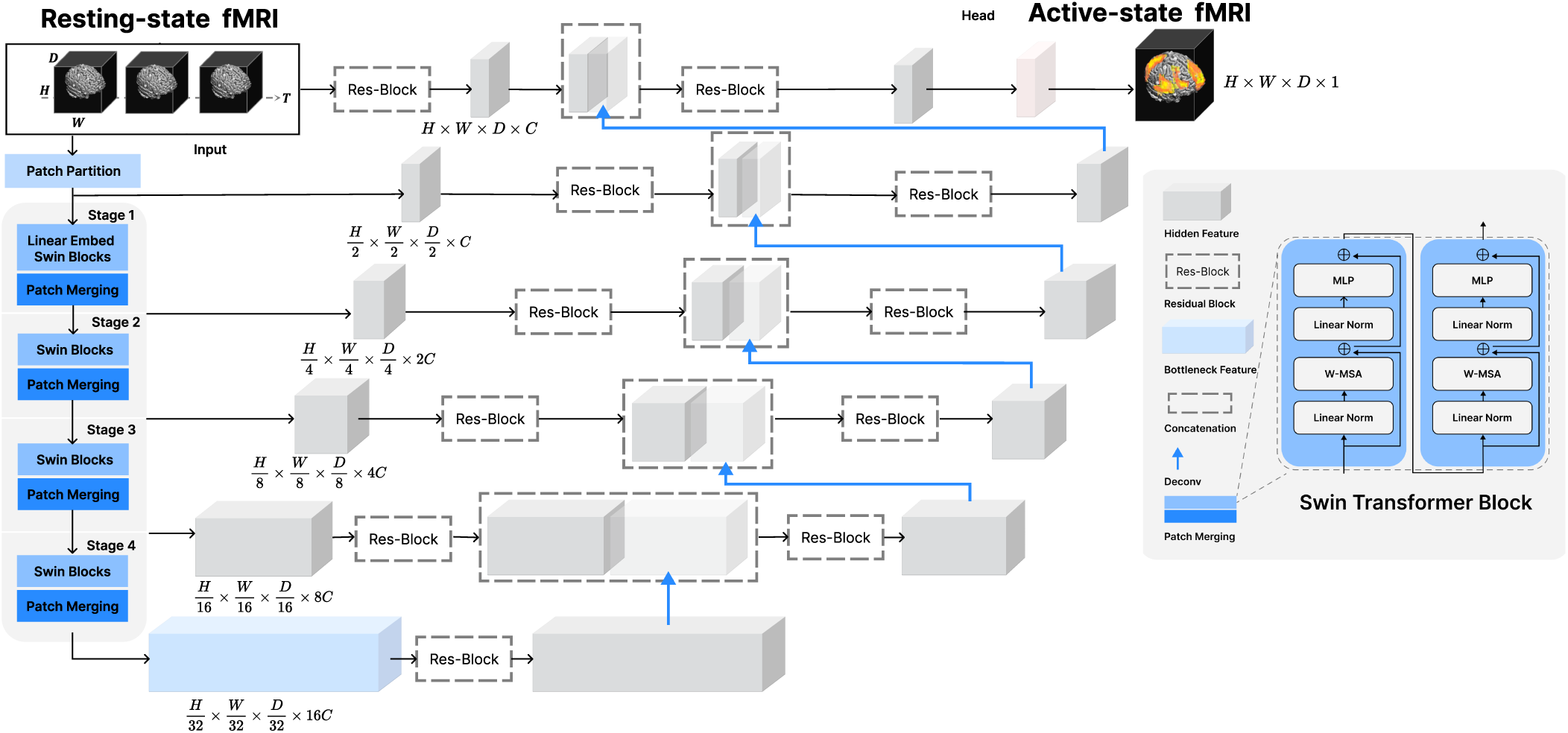
The overall architecture of Swin fMRI UNet Transformer (SwiFUN). SwiFUN takes *T* time points of resting-state fMRI volumes as input and predicts a three-dimensional task activation map. Time dimension (*T*) is considered channel dimension at the first stage. The figure is adapted from Swin UNETR (Hatamizadeh et al., 2022).

SwiFUN was trained using datasets divided into training, validation, and testing sets with a distribution ratio of 70% for training, 15% for validation, and 15% for testing. We iteratively trained the model with three different splits. We used the test set for visualization, variable prediction, and correlation analysis with head motion. To train SwiFUN, we used a Dropout rate of 0.3, initial embedding size of 24 (denotes *C* in Figure 1), and a learning rate of 5e-5 with a Cosine Annealing Warmup Restart scheduler during ten epochs. Our model was trained to minimize mean squared error (MSE), and we employed the Reconstruction-Contrastive loss as a method to improve individual identification (Ngo et al., 2022). Due to memory constraints in GPU (Nvidia A100 40GB), using an entire run of the resting state fMRI volumes (e.g., 490 volumes in UKB) as a single input was not feasible. Instead, we partitioned the fMRI volumes into sub-sequences of 30 volumes (time points) each. During training, the model learned to predict a task activation map based on these sub-sequences. When predicting a subject’s activation map for evaluation, the model first computed task activation maps for each sub-sequence, which were then averaged to produce the final task activation map for that subject. We used a mini-batch size of 4 (each containing sub-sequences) during the experiments. We also assessed the impact of the input sub-sequence length on prediction performance, as shown in Supplementary Figure S1, which depicts the effect of sequence length and mini-batch size on the performances.

### 2.3 Reconstruction-Contrastive loss

Prior studies used a contrastive loss to make the predicted activation map of a subject closely resemble their actual map while ensuring it significantly differs from the maps of other subjects (Ngo et al., 2022). We tailored this method to balance two key goals: achieving overall similarity (overall prediction accuracy) and ensuring distinct subject recognition (individual identification). In Supplementary Figure S2, we observed a trade-off between the overall prediction accuracy (measured by diagonal median) and individual identification (measured by diagonality index) during training, which supports the use of the RC (Reconstruction-Contrastive) loss.

Our loss function introduces two novel designs. Firstly, unlike previous studies that calculated the Reconstruction-Contrastive Loss *L_RC_* using just two subjects at a time (Ngo et al., 2022), we expanded this comparison to include four or more subjects. We ascertained that multiple sub-sequences of the same subject within a batch were not included in *L_C_*, ensuring that *L_C_* focuses on differences between subjects. Secondly, whereas previous methods used a two-step process of training until the same-subject error *L_R_* converges and then applying *L_C_* at certain points afterward, we introduced a new parameter *λ*, which only considers the relative weight of the two and allows for end-to-end training.

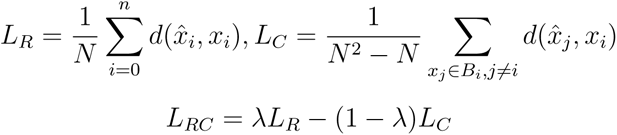

The reconstructive-contrastive loss *L_RC_* is defined as follows: Given a mini-batch of N samples B, where each sample *x_i_* represents the target 3D task activation image of a subject *i*, and *x*^ represents the corresponding prediction. *N* ^2^ − *N* in *L_C_* loss denotes the number of all possible pairs between predicted maps and actual maps from different samples in a batch. *d*() is the distance function, the mean square error (MSE) in this experiment.

### 2.4 The baseline model

#### 2.4.1 ConnTask

Several previous studies have used a GLM-based model, ConnTask, to predict task activation maps from resting state functional modes (Gal, Tik, et al., 2022; Tavor et al., 2016; Tik et al., 2021). These studies used grayordinate fMRI data (in CIFTI); however, in our study, we used the volume data (masked and converted into vectors) for a fair comparison with SwiFUN. We trained one hundred generalized linear models, each corresponding to one of the 100 cortical parcels defined in Schaefer’s atlas (Schaefer et al., 2018). These models predicted task activation maps from connectivity features (Independent Components) associated with each parcel. In the task activation maps, each region of interest (ROI) was predicted from the connectivity features of its corresponding ROI. We only used independent components as input features, treating the voxels in the connectivity features as independent training samples.

After training the models, represented by *β_k_*, we averaged *β_k_*across all subjects during the inference phase. We employed a five-fold cross-validation approach, iteratively training the models with 80% of the subjects and using the remaining 20% to predict their task activation maps. ConnTask’s five-fold cross-validation was performed within SwiFUN’s test set. Our study also explored how the number of independent components affects the predictive performance (Supplementary Table S1).

#### 2.4.2 Test-retest Contrasts

The UK Biobank (UKB) dataset includes data from repeat visits. We assessed the correlation between the task activation maps of releases 2 and 3 for the 577 participants that were present in both releases, selected from a larger pool of 7,038 datasets initially used in release 2. The correlation between the task activation maps from these two releases serves as a measure of the test-retest reliability of the actual contrast map, setting an upper bound of the model performance. The ABCD dataset includes two fMRI runs for the same emotional n-back task, both scanned on the same day. We computed a metric for test-retest reliability, similar to the UKB dataset, for the 4,934 participants who had activation maps from both runs.

### 2.5 Prediction of Individual Traits from Task Activation Maps

We evaluated the generated task activation maps by assessing how well the predicted task activation maps predicted individual traits. First, the predicted task activation maps were flattened after removing non-brain voxels using Harvard-Oxford cortical and subcortical structural atlas (Desikan et al., 2006; Frazier et al., 2005; Goldstein et al., 2007; Makris et al., 2006). The test set of SwiFUN was divided into an 80% train set and a 20% test set, and the performance was averaged over 20 iterations. To extract important features from the 132,032 valid voxels, feature reduction was performed using PCA, and the number of principal components was determined as the number with 90% explained variance. For classification and regression tasks, logistic regression with l2 regularization was used. UKB FACES, SHAPES, and FACES-SHAPES contrast were analyzed, and gender was used for the classification task, and age, mild and severe depression (given by PHQ-9 score (Manea et al., 2012)), and neuroticism (given by N-12 score, field 20127) were used for the regression task. The performance of the classification task was measured by the AUROC score between predicted and actual values, and the performance of the regression task was measured by Pearson Correlation. We conducted a permutation test to determine whether the differences between the individual trait prediction performance of ConnTask, SwiFUN with MSE and RC loss, and the actual task activation map were significant. As performed in Gal, Coldham, et al., 2022, we randomly shuffled the prediction accuracies of each model and calculated the group differences between them. The p-value was determined as the number of cases where the group difference determined by chance was higher or equal to the actual difference between the performances of two models, divided by the total number of permutations (10,000). The p-value was then Bonferroni corrected for 18 multiple comparisons, considering a total of 6 comparisons among 4 types of activation maps (real, ConnTask, SwiFUN with MSE loss and RC loss) and 3 UKB contrasts.

### 2.6 Relationship Between Head Motion and Prediction Accuracy of Task Activation Maps

We conducted an experiment to estimate the factors that can contribute to the overall quality of predicted maps. We evaluated whether the overall head motion levels of resting-state and task-state fMRI scans are correlated with the prediction accuracy of predicted maps. In the ABCD dataset, we used averaged frame-wise displacement (FD) to measure the overall head motion of resting and task-state fMRI scans. In the UKB dataset, we used the mean head motion of resting-state (field 25741) and task-related fMRI scans (field 25742) in millimeters (mm) averaged across space and time points.

## 3 Results

### 3.1. Performances Comparison in Predicting Task Activation Maps

We evaluated the efficacy of SwiFUN in predicting task activation maps from the UK Biobank (UKB) and Adolescent Brain Cognitive Development (ABCD) datasets, comparing its performance with ConnTask and test-retest reliability (Figure 2). Our analysis included a comparison between two loss functions utilized by SwiFUN: mean absolute error (MSE) and reconstructive-contrastive (RC) loss. The weight of the contrastive loss term (1 − λ) in RC loss was specified as 0.66 (refer to Supplementary Figure S3 to find the effect of the contrastive loss term). Across all contrasts within both datasets, SwiFUN showed significantly higher diagonal median performance than ConnTask (Figure 2 (a)). In the UKB data, SwiFUN’s overall accuracy was comparable to the test-retest reliability, and its accuracy surpassed the test-retest reliability of the ABCD data. SwiFUN, when trained with MSE loss, exhibited higher diagonal median performance than its RC loss counterpart in all contrasts.

**Figure 2:**
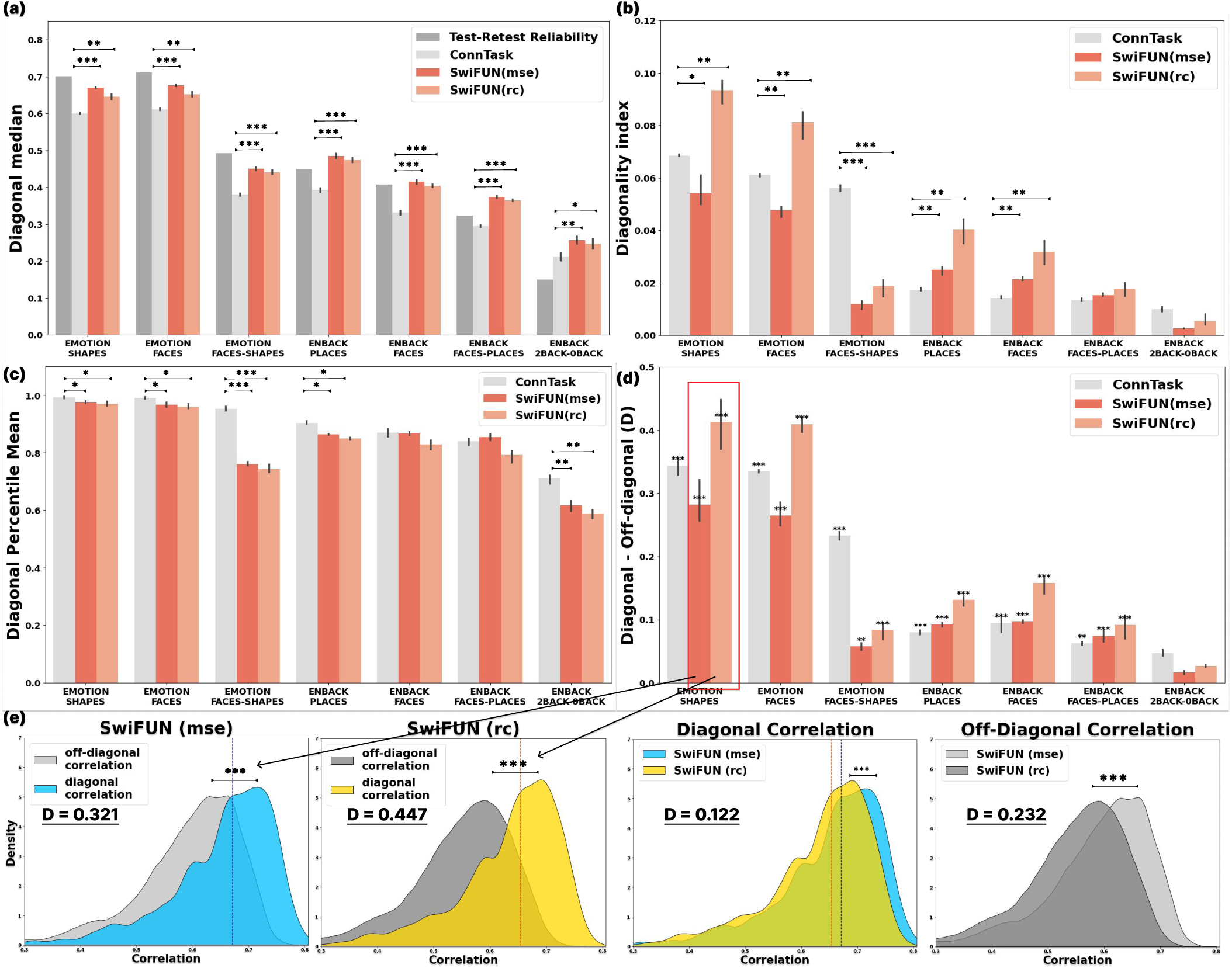
Overall performances of SwiFUN and its baseline models in UK Biobank and ABCD contrasts. (a) The diagonal Median represents the overall prediction accuracy of each model. (b) Diagonality index, (c) Diagonal Percentile Mean, and (d) Effect Size and p-value (asterisk over the bar) of the Kolmogorov-Smirnov Test (*D*) represent the model’s performance in capturing individual differences. (e) KDE plots illustrate the diagonal and off-diagonal correlations of the two kinds of SwiFUN in the UKB SHAPES Contrast. SwiFUN (mse) means SwiFUN trained with mean square error (MSE) loss, while SwiFUN (rc) denotes SwiFUN trained with Reconstruction-Contrastive (RC) loss. The dark blue and orange vertical lines mean the median of diagonal correlations of SwiFUN (mse) and SwiFUN (rc), respectively. The height of each bar represents the mean value in three repetitions, and the error bars show a 95% confidence interval. To compare two SwiFUN models and ConnTask, we conducted two-sided t-tests to assess statistically significant differences in their performance. The resulting Bonferroni-corrected p-values for three comparisons are listed in the figure (a), (b), and (c).

We then assessed the models’ ability to capture individual differences in task activation maps using a diagonality index (Figure 2 (b)). SwiFUN generally exhibited a higher diagonality index when trained with RC loss than MSE loss. Specifically, in the UKB’s SHAPES and FACES Contrast, SwiFUN trained using RC loss showed a higher diagonality index than ConnTask (*p <* 0.01), while SwiFUN trained using MSE loss did not. In ABCD’s PLACES, FACES contrasts, both types of SwiFUNs observed a higher diagonality index than ConnTask (*p <* 0.01). However, in the FACES-SHAPES contrast (UKB), SwiFUN models showed a significantly lower diagonality index than ConnTask (*p <* 0.001). In Figure 2 (c), SwiFUN with two kinds of losses showed similar diagonal percentile mean across all contrasts. ConnTask showed slightly higher (*p <* 0.05) or equal performances than SwiFUN in most contrasts except for FACES-SHAPES contrast (UKB) and 2BACK-0BACK contrast (ABCD), which shows the difference between the two conditions. Here, we could observe a similar performance drop of SwiFUNs in the FACES-SHAPES contrast as a diagonality index.

Furthermore, we conducted a Kolmogorov-Smirnov test to assess the statistical significance of the disparity between the cumulative distribution functions (CDF) of diagonal and off-diagonal correlations. In Figure 2 (d), regardless of the loss type, SwiFUN exhibited a significant distinction between diagonal and off-diagonal correlations in all task contrast maps (*p <* 0.001) except a 2BACK-0BACK contrast from ABCD data. For all contrast maps, using the RC loss led to larger effect sizes (*D*) from the Kolmogorov-Smirnov test than the MSE loss. Overall, the magnitude of the effect size (*D*) was similar to that of the diagonality index. Figure 2 (e) shows why SwiFUN trained with RC loss has a higher effect size (*D*) in the Kormogorov-Smirnov test than the model trained with MSE loss. In UKB SHAPE contrast, the diagonal correlations of MSE loss were significantly lower than RC loss’s (*D* = 0.122, *p <* 0.001), but the off-diagonal correlations of RC loss were much lower overall than MSE loss’ (*D* = 0.232, *p <* 0.001). This means that RC loss maximizes the individual uniqueness of predicted task contrast maps by decreasing off-diagonal correlations to a greater extent than the decrease in diagonal correlations, resulting in an overall improvement in the specificity of the predicted maps.

### 3.1 Qualitative Evaluation on Volumetric Task Activation Map

Figure 3 display FACES contrast maps in the emotional matching task (UKB) and emotional n-back task (ABCD), showing group-averaged task activation map and individual maps of three subjects with the top-1, top 25%, and top 50% correlation between the actual and predicted maps using SwiFUN with MSE loss. For the FACES contrasts in both datasets, SwiFUN successfully predicted the activations observed in regions related to face recognition, such as the Occipital lobe, Fusiform Face Area, and Amygdala. Additionally, as the white arrows show, SwiFUN effectively captured the individual differences within the brain regions.

**Figure 3:**
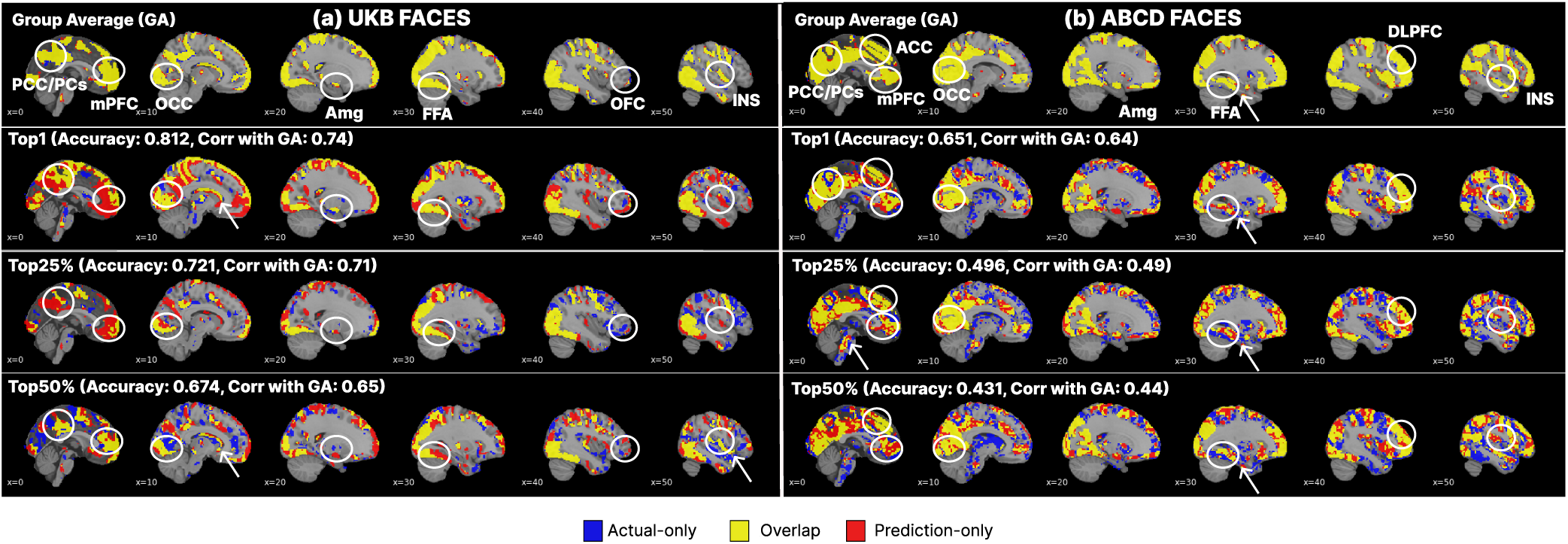
FACES contrast maps from UKB and ABCD task-based fMRI data in a series of sagittal views estimated by SwiFUN. Voxel values were normalized using z-scoring and then thresholded at the 97th percentile (two-sided). The yellow regions are cases where the areas predicted by SwiFUN match the actual activation, the blue regions are actual activations that SwiFUN did not predict, and the red regions are cases where SwiFUN predicted activation but no actual activation occurred. Accuracy indicates the correlation between the predicted and actual task activation map, and Correlation with GA means the correlation between the group-averaged map and the actual map. The brain voxels in the Harvard-Oxford atlas were used for visualization. White circles indicate brain regions that were prominently shown in the group-averaged maps, and white arrows represent the brain regions showing individual differences captured by SwiFUN. Abbreviations of brain regions are as follows: Posterior cingulate cortex (PCC), Medial prefrontal cortex (mPFC), Amygdala (Amg), Fusiform area(FFA), Orbitofrontal cortex (OFC), Insula (INS), Dorsolateral prefrontal cortex (DLPFC).

The average of the actual task activation maps and the average of the task activation maps predicted by SwiFUN were highly consistent (*r*=0.99 for both contrasts). However, while the actual activation maps showed low similarity to the representative group-averaged maps (average r=0.625 for UKB FACES and r=0.4 for ABCD FACES contrasts), the predicted activation maps exhibited patterns similar to the group-averaged maps (average r=0.949 for UK Biobank data and r=0.96 for ABCD data). Therefore, the more similar the actual activation maps were to the representative activations, the more accurately SwiFUN tended to predict those activation maps (r=0.981 in UK Biobank data and r=0.991 in ABCD data).

### 3.3 Head Motion is negatively correlated with Prediction Accuracy of SwiFUN

We verified that transient noise sources such as head motion in fMRI data are significantly associated with how well the task activation map reflects representative activation and the predictive model’s accuracy. Figure 4 shows that SwiFUN’s prediction accuracy, as measured by the diagonal median, is significantly and negatively correlated with head motion level (*P <* 0.001). In both the UK Biobank and ABCD datasets, the mean head motion level in task fMRIs (overall mean of -0.391) was more negatively correlated with the model’s performance than the mean head motion level in resting-state fMRIs (overall mean of -0.2377). We found the difference in correlation coefficient between task fMRI and resting-state fMRI data was significant in all contrasts through Fisher’s Z-Transformation (*P <* 0.001). In addition, the Pearson correlation between head motion and prediction accuracy in the contrasts featuring overall brain activity (i.e., FACES, SHAPES, and PLACES) (mean of -0.3322) was stronger compared to the negative correlation (mean of -0.2905) in the region-specific contrasts (i.e., FACES-SHAPES, 2BACK-0BACK). The correlation between SwiFUN’s prediction accuracy and head motion showed an overall stronger negative correlation compared to the corresponding correlation in the ConnTask, but the difference between the two predictive models was not significant (Supplementary Figure S4). we also found that the higher each subject’s head motion, the lower the similarity between the actual task activation map and the group-averaged map. (Supplementary Figure S5)

**Figure 4:**
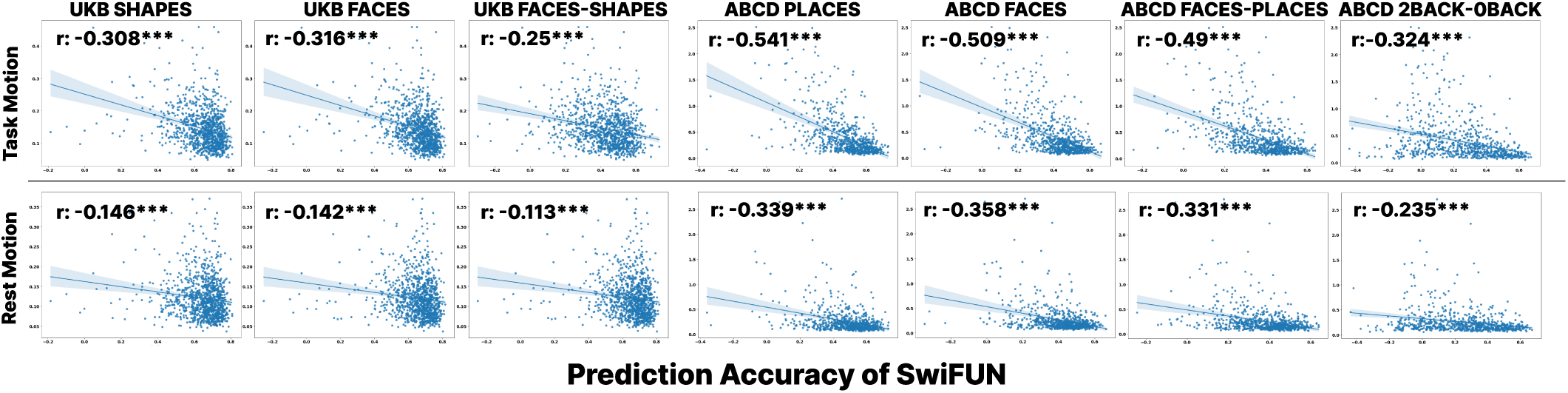
Scatter plots showing a negative correlation between mean head motion and prediction accuracy of SwiFUN. Each row represents the averaged head motion of task-based fMRI and resting-state fMRI. Prediction Accuracy means the correlation between the predicted and actual task activation map.

### 3.4 Prediction of Individual Traits from the Predicted Task Activation Maps

We evaluated how well task activation maps predicted by predictive models captured individual differences by assessing their accuracy in predicting sex, age, depression severity (measured by PHQ-9 scores), and neuroticism levels (measured by N-12 scores) from task activation maps derived from the UKB data. From Figure 5 (a) and (b), the SwiFUN models trained with MSE and RC loss outperformed both the actual task activation maps and the maps predicted by ConnTask in identifying sex and age across all contrasts (*p <* 0.001). For predicting depressive symptoms, as shown in Figure 5 (c), activation maps predicted by SwiFUN models showed better performance for the SHAPES (average *r* = 0.153) and FACES (average *r* = 0.119) contrasts compared to the FACES-SHAPES contrast (*r* = −0.012). The activation maps predicted by SwiFUN models trained with RC loss led to higher Pearson correlation scores for predicting depression than the ConnTask-predicted maps across all contrasts (*p <* 0.001), as well as higher scores than the actual maps for the SHAPES (*p <* 0.05) and FACES-SHAPES (*p <* 0.01) contrasts. For predicting neuroticism scores, SwiFUN models outperformed ConnTask across all contrasts (*p <* 0.05). However, compared to actual task activation maps, SwiFUN showed significantly better predictive performance only for the FACES contrast (*p <* 0.01). While ConnTask outperformed actual maps in predicting sex (*p <* 0.001), it exhibited comparable or lower prediction performances than actual maps for age, depression, and neuroticism. The performance differences between SwiFUN models trained with mean squared error (MSE) loss and RC loss were insignificant across all variables and contrasts.

**Figure 5:**
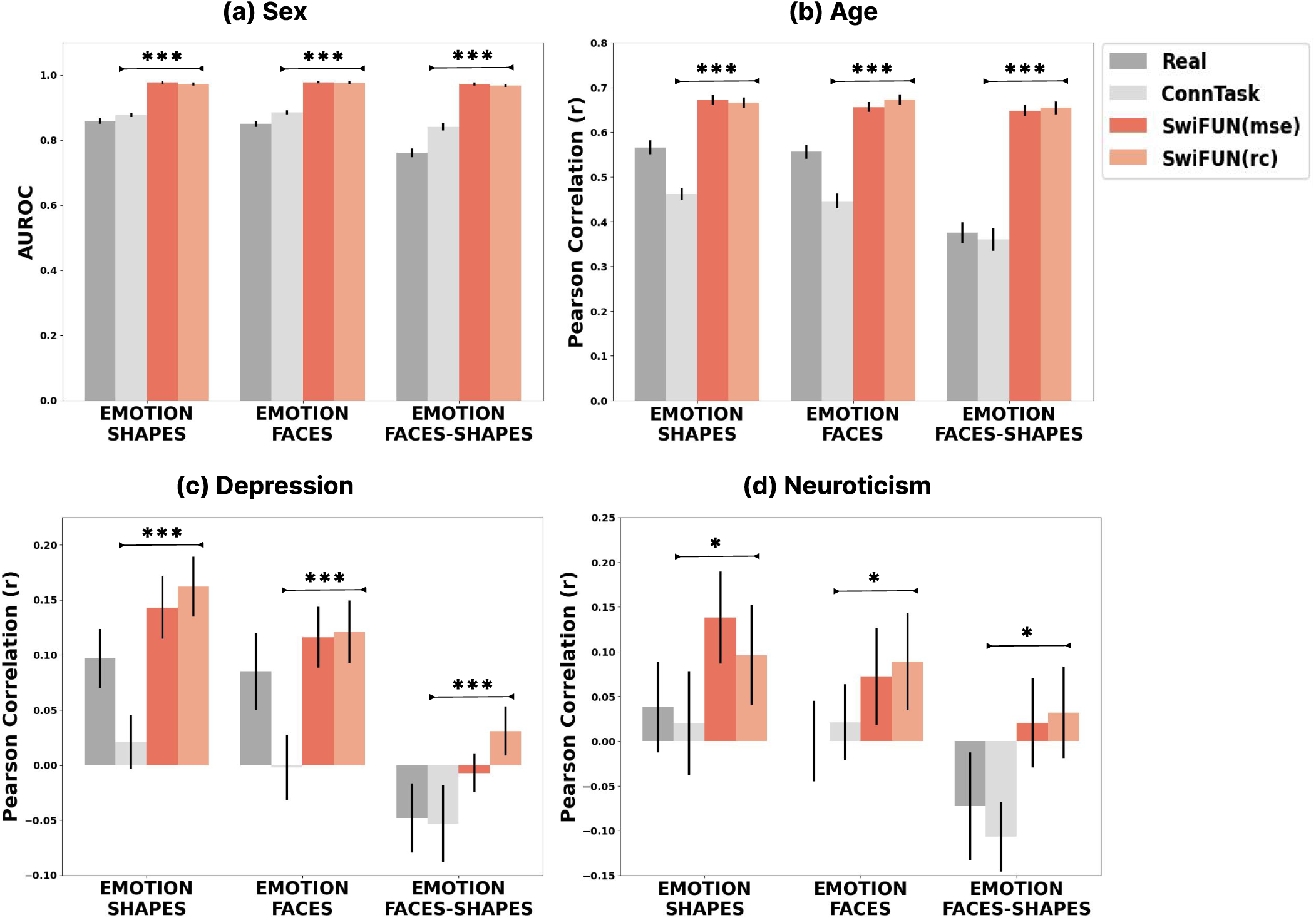
Predictive performance of actual and predicted task activation maps for individual traits (sex, age, depression, and neuroticism) using UK Biobank data. The height of each bar represents the mean value in twenty repetitions, and the error bars show a 95% confidence interval.

## 4 Discussion

In this study, we introduce SwiFUN, a novel deep neural network that merges the fMRI Transformer’s capacity to learn spatiotemporal patterns from resting-state fMRI data with the predictive capabilities of the U-Net architecture. This combination enables the accurate prediction of task-specific brain activations. Unlike previous models that utilized surface fMRI data through complex processing pipelines, SwiFUN employs a more universally applicable end-to-end deep learning approach with volumetric fMRI data. By leveraging resting-state and task-related fMRI data from the ABCD and UKB datasets, SwiFUN outperforms existing GLM-based models in predicting task-related brain activity and capturing individual differences. Furthermore, the predicted task activation map can further predict an individual’s biological and psychological traits. Our approach offers new possibilities for studying brain (dys-)function without relying on laborious feature engineering processes.

SwiFUN exhibits the potential to predict functional activation patterns with high accuracy. In adults, the UKB data, the overall accuracy of SwiFUN was comparable to the test-retest reliability, indicating that the model’s performance was on par with the inherent stability of the functional data. In children, the ABCD data, SwiFUN demonstrated higher accuracy than the test-retest reliability. This unexpected finding may be related to the challenges present in ABCD task fMRI data: e.g., suboptimal experimental designs, head movement, and decreased attention in youth (Casey et al., 2018). These factors can lead to noisy and lower-quality task activation maps. SwiFUN’s superior performance suggests that it may be particularly valuable for inferring functional activation patterns in datasets with suboptimal data quality, including those involving clinical populations or pediatric samples.

Consistent with previous literature (Zheng et al., 2022), all the predictive models demonstrated lower individual identification performance for contrasts reflecting the difference between two conditions (e.g., FACES-SHAPES, FACES-PLACES, 2BACK-0BACK) compared to single-condition contrasts (e.g., FACES, SHAPES) (Figure 2). This gap was more pronounced for the SwiFUN model than for ConnTask. One possible explanation might be the differences in how these two models leverage resting-state fMRI information to predict task-based activations. While ConnTask utilizes only the functional networks corresponding to each seed region (through ICA modeling) as the input, SwiFUN uses the spatiotemporal relationships among the entire brain regions (through shifted-window multi-head self-attention). For the contrasts that broadly capture the activity of the widespread brain regions, SwiFUN may have advantages; however, for the contrasts recruiting more focal brain activity (differences) (e.g., amygdalar activity differences in FACES-SHAPES), the additional information from the whole-brain resting-state data may not confer the same benefit.

We observed the characteristics of task activation maps that are less similar to the group-averaged map and their impact on SwiFUN’s prediction accuracy. In Figure 3, SwiFUN tended to predict the task activation maps less accurately when the task activation maps are unique, which are dissimilar to the group-averaged map. The more unique task activation maps were, the higher the head motion level their original resting-state or task-based fMRI had, as demonstrated in Supplementary Figure S5. Furthermore, we found that the level of head motion in the task fMRI data, which corresponds to the ground truth of the prediction, has a much more negative impact on the model’s prediction performance than the head motion in the input resting-state fMRI data. This relationship between head motion and the model’s predictive performance suggests that the reason for the model’s low prediction performance for certain subjects is transient factors like the low quality of the actual activation maps.

Our results underscore SwiFUN’s capacity to elucidate individual characteristics, including depression, neuroticism, sex, and age. This is consistent with previous findings that task activation maps predicted by resting-state fMRI better predict cognitive variables such as intelligence than real task activation maps (Gal, Coldham, et al., 2022; Gal, Tik, et al., 2022). This advancement suggests that resting-state fMRI-derived task activation maps, particularly those generated by SwiFUN, hold significant potential to reflect cognitive and biological traits more accurately than traditional task activation maps, which may be under the influence of nuisance variables (head motion, scanning artifacts, attention level fluctuations) (Bernstein-Eliav & Tavor, 2022). Such capability implies broader applications for SwiFUN, including the potential for diagnosing and predicting psychiatric disorders, thereby positioning it as a valuable tool in neuroscientific research and clinical practice. Additionally, SwiFUN’s framework, leveraging an attention mechanism-based deep neural network, is well-equipped to integrate additional imaging modalities, like T1-weighted structural MRI or diffusion-weighted MRI. This capability may not only enrich its application in neuroimaging analysis but specifically bolster its potential for more comprehensive and nuanced modeling of brain function and dysfunction.

Naturalistic fMRI, such as using movies as stimuli, offers a powerful lens for studying brain function in real-world contexts, potentially outperforming traditional methods (resting-state fMRI or task-based fMRI) in predicting individual traits and mapping brain activity (Bernstein-Eliav & Tavor, 2022; Finn & Bandettini, 2021; Gal, Coldham, et al., 2022). However, the dynamic nature of naturalistic stimuli poses challenges for conventional fMRI analysis techniques. SwiFUN, with its ability to capture spatiotemporal changes in brain activity, may emerge as a promising solution for unlocking the rich information embedded within naturalistic fMRI data.

Our study faces three primary limitations. First, understanding which brain regions significantly affect the prediction of task activation maps is a challenge. Previous research has linked task-related brain activity with specific resting-state functional connectivity using generalized linear models (Izakson et al., 2023; Tik et al., 2021). However, SwiFUN’s use of multiple nonlinear transformations complicates the clarity of these input-output relationships. In fMRI studies utilizing deep neural networks, Explainable AI (XAI) methods such as Grad-CAM and Integrated Gradients are widely used to pinpoint brain areas relevant to the single target outcome (Kim et al., 2024; S. Nguyen et al., 2020). However, these techniques are not yet adept at dissecting spatiotemporal features of four-dimensional resting-state fMRI inputs attributing to the resultant three-dimensional task-related activation maps. Thus, creating effective interpretation methods for SwiFUN remains a crucial future objective. Second, the current model cannot process the entire fMRI sequence at once due to hardware limitations, with our Nvidia A100 40GB GPU managing only up to 30 volumes simultaneously. This limitation hinders the model’s ability to capture extended brain dynamics, which can last for significant periods. Enhancing the model to accommodate longer input sequences represents a vital future goal. Lastly, a separate model must be trained for every distinct task and condition, which is not only resource-intensive but also overlooks potential commonalities across various tasks and conditions (Ngo et al., 2022). M. Nguyen et al., 2023 has recently shown that providing resting-state functional connectivity and group-activation maps as inputs to deep learning models can enable the prediction of individualized activation maps. Likewise, developing learning strategies that can be applied to diverse tasks and conditions through a single training process is a crucial challenge for future research.

## 5 Conclusion

This study suggests that training deep neural networks that capture spatiotemporal patterns directly from fMRI data may contribute to better performance in task activation prediction. In the future, we anticipate predicting various task-related brain activity from just a few minutes of resting-state fMRI data, significantly reducing the scanning time and effort required to capture task-based fMRI.

## Ethics Statement

The research presented in this work analyzes functional magnetic resonance imaging (fMRI) data obtained from individuals who participated in the Adolescent Brain Cognitive Development (ABCD) study and the UK Biobank project. Participation in these studies was voluntary, and all individuals provided written consent after being informed about the studies’ protocols and procedures in compliance with the ethical standards and guidelines established for each respective study.

## Data and Code Availability

The access procedures of the Adolescent Brain Cognitive Development (ABCD) study and UK Biobank specify that only approved researchers can access participant data. Therefore, the data from this study cannot be publicly released. The code for using SwiFUN is available at https://github.com/Transconnectome/SwiFUN. The code to run our baseline ConnTask is available at https://github.com/ShacharGal/connTask.

## Author Contributions

Junbeom Kwon: Junbeom Kwon served as the primary investigator and was instrumental in the development and conceptualization of the model. Junbeom Kwon led the experimental design, conducted the majority of the experiments, and was responsible for establishing the experimental setup. Junbeom Kwon also led data analysis and interpretation and drafted the original manuscript.

Jungwoo Seo: Jungwoo Seo contributed by running the baseline models, executing the initial data preprocessing and first-level analysis, and assisting with drafting and revising the manuscript. Additionally, Jungwoo Seo reviewed the literature, integrating relevant findings into the study. He also played a key role in discussing the relationship among transient factors and their impact on model performance.

Heehwan Wang: Heehwan Wang provided critical insights and discussions that helped in the conceptual framework and refinement of the model. Heehwan Wang also contributed to the review and editing of the manuscript, enhancing its overall quality and coherence.

Taesup Moon: Taesup Moon provided essential technical support and advised on technical preciseness, ensuring the research methodology adhered to the highest scientific standards.

Shinjae Yoo: In addition to providing technical support and advice, Shinjae Yoo also provided access to advanced computational resources at NERSC Perlmutter, which were crucial for the project’s data processing and analysis phases.

Jiook Cha: Jiook Cha, as the corresponding author, took on a supervisory role, providing overall guidance and direction for the project. Jiook Cha was pivotal in securing funding and resources, ensuring the project’s success.

## Supplementary Material

### 1. Effect of input sequence length and batch size on the predictive performance of SwiFUN

In Figure S1 (a), we confirmed whether the sequence length impacts the overall concordance (diagonal median) and individual identification (diagonality) of SwiFUN. SwiFUN was trained on the SHAPES contrast map using MSE loss. The experiments were conducted with a fixed mini-batch size of 4, and four different settings were tested with lengths of 10, 20, 30, and 48 considering the limitations of GPU memory (Nvidia A100 40GB GPU). The overall performance differences were not substantial, but the diagonal median showed improvement as the sequence length increased, reaching its peak at length 30. On the other hand, the diagonality index displayed the best performance at length 30 but did not exhibit consistent improvement with increasing length. Considering these results and the training speed associated with sequence length, all experiments were conducted with a length of 30.

Furthermore, we confirmed whether SwiFUN trained with RC loss is impacted by the mini-batch size (Figure S1 (b)). Considering that the contrastive loss compares samples within the mini-batch, we hypothesized that the mini-batch size would impact performance. In the RC loss, we set the value of 1−λ (the weight of the contrastive loss term) to 0.33. Our analysis revealed that in all tasks, the diagonal median exhibited a small increase as the batch size increased. On the other hand, the diagonality index showed a decrease with larger batch sizes. This suggests that increasing the number of samples in a batch can weaken the effect of contrastive loss.

**Figure S1:**
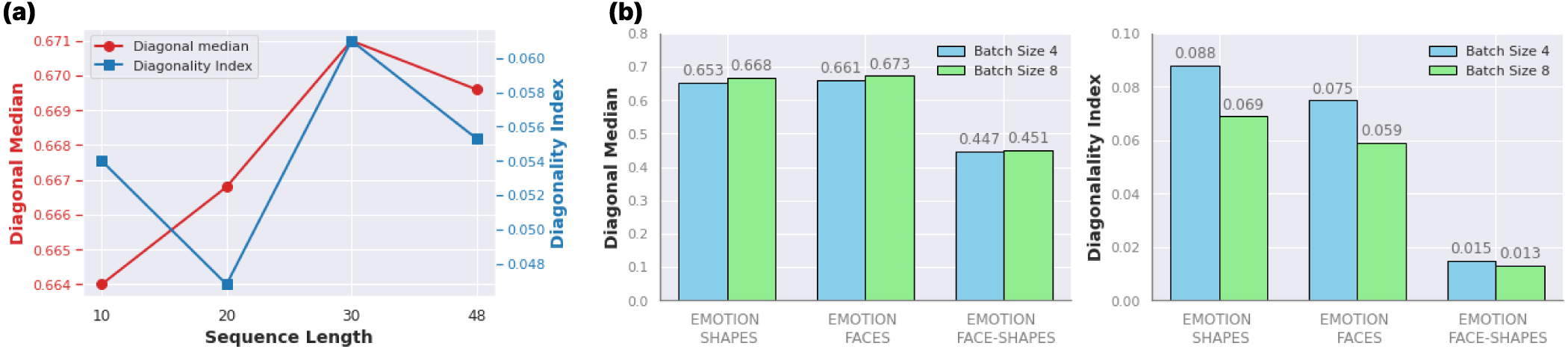
Effect of sequence length and batch size on the performance of SwiFUN

### 2. Trade-off between overall prediction accuracy and individual identification

As shown in Figure S2, during the training process of SwiFUN with MSE loss, the diagonal median and diagonality index initially increase together, but at some point, the diagonal mean starts decreasing while the diagonality index continues to increase. This indicates that initially, the model is trained to increase overall prediction accuracy, similar to the group mean activity. However, at a certain point, the model shifts its focus towards capturing subtle individual differences at the expense of overall prediction accuracy. However, there is a drawback regarding the sharp decrease in the diagonal median compared to the increase in the diagonality index. Therefore, in this study, we addressed this issue by incorporating the Reconstruction-Contrastive loss, allowing the model to train in a direction that avoids excessive convergence towards group means and instead reveals individual differences.

**Figure S2:**
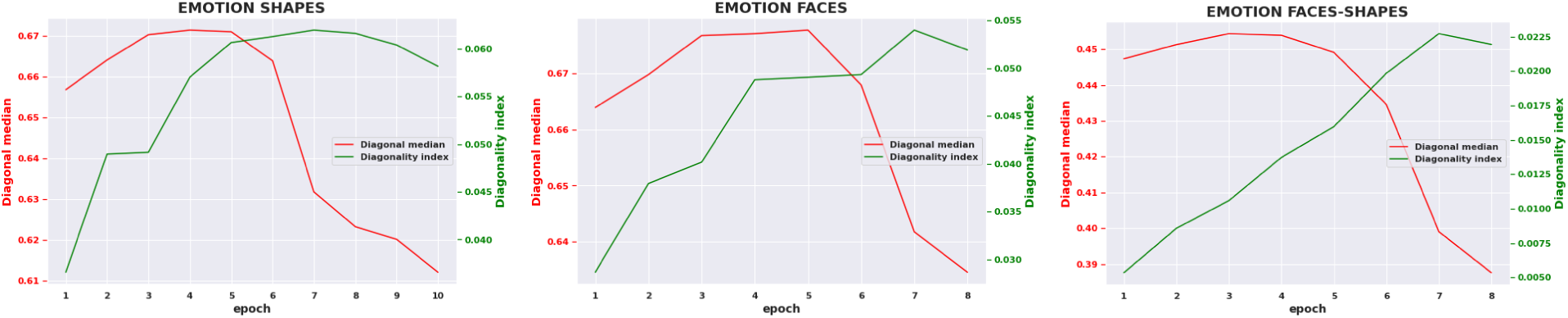
Training Curve of Diagonal Mean and Diagonality Index

### 3. Performance of ConnTask with different numbers of independent components

In Table S1, we compared the performances of ConnTask with different independent components(IC) over three emotion contrasts in UKB data. We observed that more number components (55) have a positive effect on both overall prediction accuracy (diagonal median) and individual identification (diagonality index, Effect size *D* of Kolmogorov-Smirnov test, and Diagonal Percentile Mean) than fewer components (21). Thus, we compared ConTasks, which used 55 independent components, with SwiFUN.

**Table S1:** Performance of ConnTask.

| Task | IC | Diagonal Median | Diagonality Index | $KS\_D$ | Diag. Percent. Mean |
| --- | --- | --- | --- | --- | --- |
| SHAPES | 21 | $0.593 \pm 0.002$ | $0.062 \pm 0.000$ | $0.318 \pm 0.015$ | $0.991 \pm 0.003$ |
| SHAPES | **55** | $0.600 \pm 0.001$ | $0.069 \pm 0.000$ | $0.344 \pm 0.013$ | $0.993 \pm 0.003$ |
| FACES | 21 | $0.604 \pm 0.001$ | $0.055 \pm 0.000$ | $0.305 \pm 0.001$ | $0.989 \pm 0.003$ |
| FACES | **55** | $0.611 \pm 0.002$ | $0.061 \pm 0.001$ | $0.335 \pm 0.002$ | $0.992 \pm 0.004$ |
| F-S | 21 | $0.371 \pm 0.004$ | $0.05 \pm 0.001$ | $0.207 \pm 0.005$ | $0.943 \pm 0.01$ |
| F-S | **55** | $0.380 \pm 0.003$ | $0.056 \pm 0.001$ | $0.233 \pm 0.006$ | $0.953 \pm 0.008$ |

### 4. The effect of contrastive loss term in Reconstruction-Contrastive loss

In Figure S3, we investigated how the diagonal median and diagonality index vary with the adjustment of the weight of the contrastive loss in the RC loss. We conducted experiments with three settings for 1 − λ: 0 (*L_R_* only), 0.5 (*L_R_* : *L_C_* = 1 : 1), 0.6 (*L_R_* : *L_C_* = 1 : 1.5), and 0.66 (*L_R_* : *L_C_* = 1 : 2). The results showed that as we increased the weight of the contrastive loss term in all contrast maps, the diagonal median decreased while the diagonality index increased. However, compared with the results of predicting the SHAPES contrast map in Figure 5, where no contrastive loss term was used, we can observe that the increase in the diagonality index is much more significant compared to the relatively small decrease in the diagonal median. For instance, in the case of predicting the SHAPES contrast in Figure 5, after the diagonal median converged at 0.671, it decreased by 0.04, while the diagonality index increased by only 0.001. On the other hand, while 1 − λ increased from 0 to 0.66, the diagonality index significantly improved by 0.027, with a similar decrease in the diagonal median by 0.05. This indicates that by using the RC loss, it is possible to effectively enhance individual identification performance while sacrificing overall concordance to a lesser extent.

**Figure S3:**
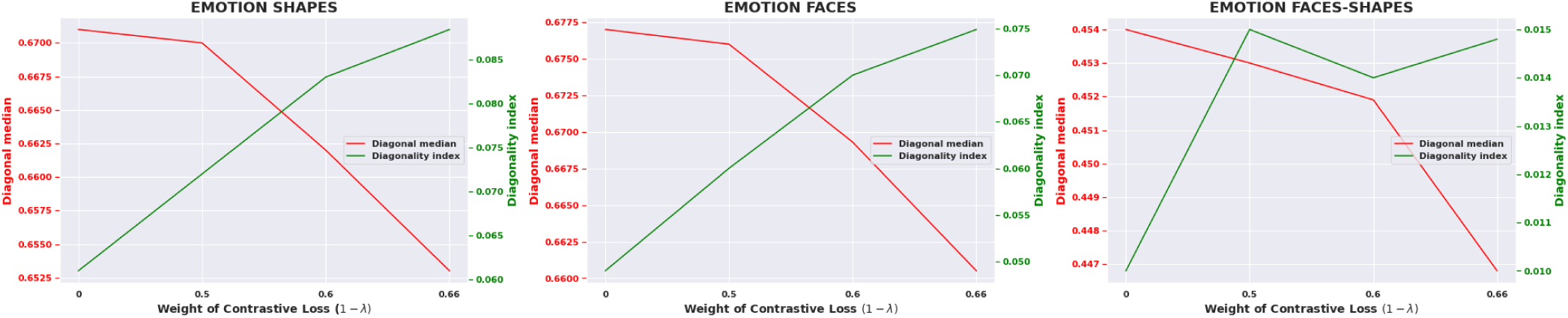
Effect of Contrastive loss term in Reconstruction-Contrastive loss

### 5. Negative Correlation between Head Motion Level of fMRI Scans and ConnTask’s Prediction Performance

**Figure S4:**
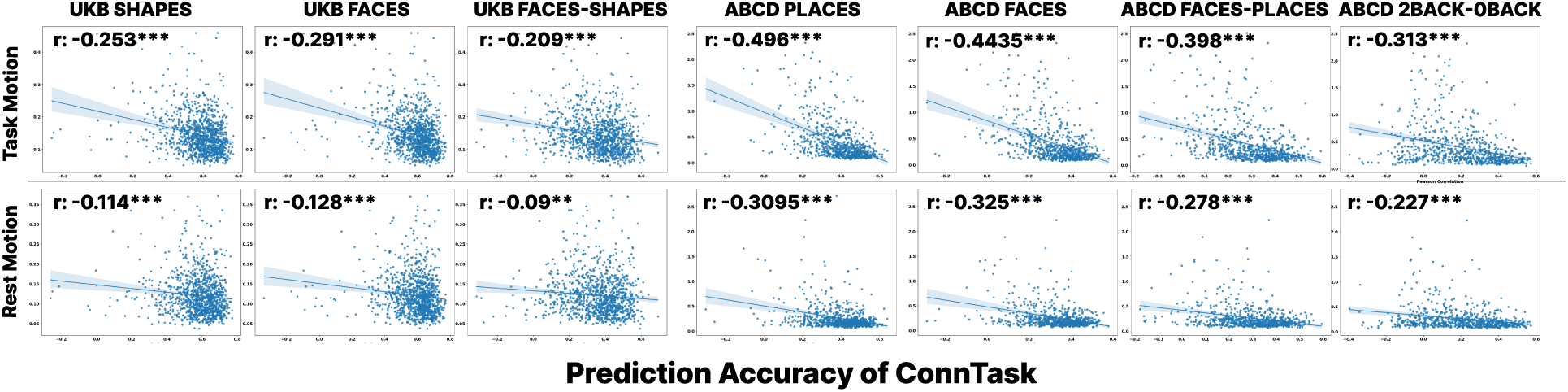
Scatter plots showing a negative correlation between mean head motion and prediction accuracy of ConnTask. Each row represents the averaged head motion of task-based fMRI and resting-state fMRI.

### 6. Negative Correlation between Head Motion Level of fMRI Scans and Similarity between the Actual Map and Group Average Map

**Figure S5:**
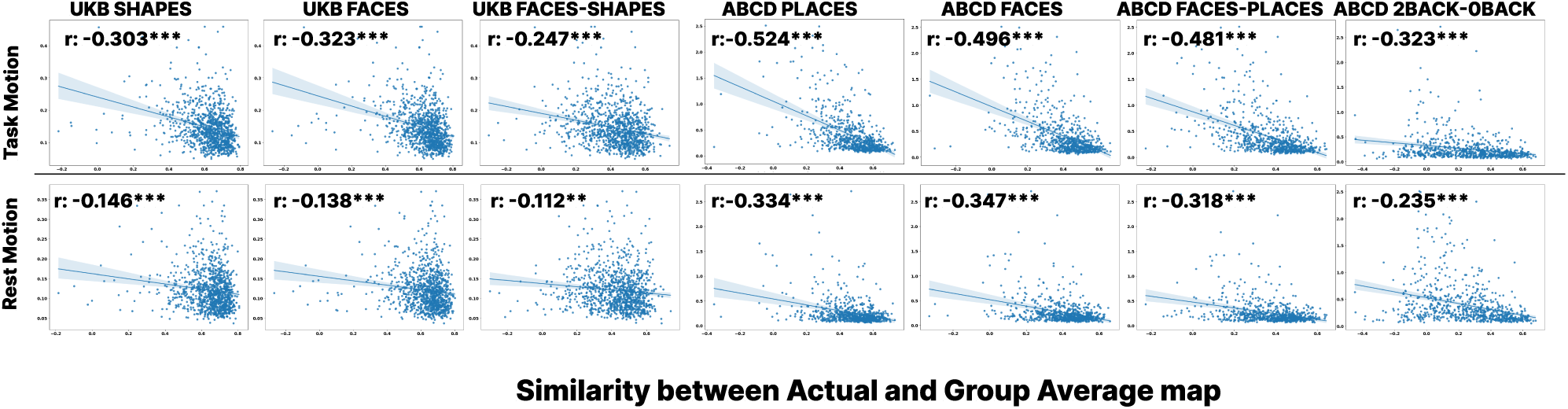
Scatter plots showing a negative correlation between mean head motion and group average map. Each row represents the averaged head motion of task-based fMRI and resting-state fMRI.

